# Cortical timescales and the modular organization of structural and functional brain networks

**DOI:** 10.1101/2023.07.12.548751

**Authors:** Daniel J. Lurie, Ioannis Pappas, Mark D’Esposito

## Abstract

Recent years have seen growing interest in characterizing the properties of regional brain dynamics and their relationship to other features of brain structure and function. In particular, multiple studies have observed regional differences in the “timescale” over which activity fluctuates during periods of quiet rest. In the cerebral cortex, these timescales have been associated with both local circuit properties as well as patterns of inter-regional connectivity, including the extent to which each region exhibits widespread connectivity to other brain areas. In the current study, we build on prior observations of an association between connectivity and dynamics in the cerebral cortex by investigating the relationship between BOLD fMRI timescales and the modular organization of structural and functional brain networks. We characterize network community structure across multiple scales and find that longer timescales are associated with greater within-community functional connectivity and diverse structural connectivity. We also replicate prior observations of a positive correlation between timescales and structural connectivity degree. Finally, we find evidence for preferential functional connectivity between cortical areas with similar timescales. We replicate these findings in an independent dataset. These results contribute to our understanding of functional brain organization and structure-function relationships in the human brain, and support the notion that regional differences in cortical dynamics may in part reflect the topological role of each region within macroscale brain networks.

## Introduction

Even at rest, brain activity is in constant flux. Studying the properties of these fluctuations and their relationship to other aspects of brain structure and function can provide important insights into functional brain organization. There are many different ways to characterize the properties of neural time series, ranging from simple, commonly used metrics such as amplitude and frequency, to complex, seemingly esoteric measures taken from physics and and the study of dynamical systems (Breakspear, 2017; Fulcher et al., 2013; Fulcher & Jones, 2017; Lubba et al., 2019; Toker et al., 2020). From this menagerie of measures, the past decade has seen estimates of neural “timescale” emerge as an increasingly popular way to study the dynamical properties of brain activity. While there are multiple ways to estimate the timescale of neural activity, their common goal is to characterize the degree to which brain activity (or the information it represents) is similar from one moment to the next. The literature on neural timescales ranges from studies using invasive recordings of individual neurons and population activity in animals (Cirillo et al., 2018; Maisson et al., 2021; Murray et al., 2014; Nougaret et al., 2021; Rossi-Pool et al., 2021; Runyan et al., 2017; Wasmuht et al., 2018) to BOLD fMRI and MEG/EEG in humans (Baldassano et al., 2017; Fallon et al., 2020; Golesorkhi et al., 2021; Hasson et al., 2008; Huang et al., 2018; Ito et al., 2020; Müller et al., 2020; Raut et al., 2020; Stephens et al., 2013; Watanabe et al., 2019; Wengler et al., 2020; Zilio et al., 2021), as well as computational models which seek to provide mechanistic explanations for regional differences in timescale (Chaudhuri et al., 2014, 2015; Demirtaş et al., 2019; Gollo et al., 2015; Li & Wang, 2022). Many of these studies suggest the presence of a “hierarchy of timescales” in the cerebral cortex which parallels information processing hierarchies, with shorter timescales in sensory and motor areas responsible for processing rapidly changing low-level sensory information and action sequences, and longer timescales in higher-order association regions which integrate information over longer periods in order to support more complex information processing (such as the construction and maintenance of increasingly abstract representations).

Computational models and emerging empirical work suggest that regional differences in neural timescales are shaped not only by heterogeneity in local circuit properties, but also by the pattern of connections each region receives from distant brain areas (Baria et al., 2013; Chaudhuri et al., 2014, 2015; Demirtaş et al., 2019; Fallon et al., 2020; Gao et al., 2020; Gollo et al., 2015; Ito et al., 2020; Li & Wang, 2022; Müller et al., 2020; Sethi et al., 2017; Watanabe et al., 2019; Wengler et al., 2020). In particular, emerging evidence suggests that neural timescales may systematically vary between brain areas with different topological properties, with highly connected (i.e. high degree) “hub” regions tending to exhibit longer timescales than less topologically central areas (Baria et al., 2013; Fallon et al., 2020; Gollo et al., 2015; Sethi et al., 2017).

However, while degree is arguably the most fundamental measure of regional topology (and the network feature which has most consistently been associated with regional dynamics in prior studies), it fails to consider another key aspect of brain network topology. In particular, in addition to high degree hubs, an extensive literature in network neuroscience has demonstrated that brain networks exhibit a modular community structure in which regions are more likely to connect to (or interact with) other regions in the same community than to regions in other communities (Girvan & Newman, 2002; Newman, 2006; Sporns & Betzel, 2016). This modular structure can be found at multiple scales, ranging from microscale neuronal populations (Lee et al., 2016; Schroeter et al., 2015; Shimono & Beggs, 2015) to macro-scale functional and structural brain networks (Crossley et al., 2013; Hagmann et al., 2008; He et al., 2009; Meunier et al., 2009; Power et al., 2011). This architecture is thought to play a key role in facilitating functional specialization and flexible reconfiguration by balancing the competing demands of integration and segregation (Bertolero et al., 2015, 2018; Bullmore & Sporns, 2009; Sporns, 2013; Sporns & Betzel, 2016). By considering this modular organization, we can take a much more nuanced view of the role each region plays within the larger network. In particular, beyond identifying “global” hubs (as has been done in prior studies of the relationship between timescales and network topology), we can characterize the degree to which each brain area exhibits widespread connectivity within its own community, and/or diverse connectivity to regions in other communities (Guimerà & Amaral, 2005; Guimerà & Nunes Amaral, 2005; Power et al., 2013; van den Heuvel & Sporns, 2013a).

We hypothesized that regional differences in intrinsic brain dynamics would be associated with the modular organization of human brain networks, and specifically that brain areas with extensive within-community connectivity (i.e. high within-module degree) or diverse connectivity across communities (i.e. high participation coefficient) would tend to have longer timescales, consistent with their putative roles in integrating information within and between network modules. To test these hypotheses, we mapped cortical timescales in human participants using resting BOLD fMRI, and characterized the multi-scale community structure of functional and structural brain networks. Our key methods and primary hypotheses were preregistered via the Open Science Framework (Lurie, 2018), and we provide an internal replication of all analyses in a second independent dataset.

## Methods

### Resting BOLD fMRI data

For our initial (“discovery”) analysis, we analyzed data from 93 healthy young individuals (ages 18-35) from the Enhanced Nathan Kline Institute-Rockland Sample (Nooner et al., 2012). These individuals were selected based on the absence of any history of psychiatric or neurological illness, as well as the presence of complete imaging data (i.e. no missing scans). Resting fMRI scans were 10 minutes in duration with TR/TE = 1400/30ms and 2mm isotropic voxels. Data were preprocessed using FMRIPRPEP (Esteban et al., 2019). GLM denoising was applied using Nilearn (Abraham et al., 2014) and included 24 motion parameters plus 6 aCompCor components containing nuisance signals from WM and CSF voxels (Behzadi et al., 2007), global signal regression, and a 0.009 to 0.08 Hz bandpass filter.

For our validation analysis, we selected a sample of 100 unrelated healthy young individuals (ages 21-35) from the Human Connectome Project (Van Essen et al., 2013) based on a subject list provided by (Shine et al., 2016). For each individual, we analyzed the first 15 minute resting fMRI scan (TR/TE = 720/33ms, 2mm isotropic voxels) from the first scan session.. We used minimally preprocessed data (Glasser et al., 2013) provided by HCP which had undergone ICA-FIX denoising (Griffanti et al., 2014), global signal regression, and a 0.009 to 0.08 Hz bandpass filter.

Astute readers may note that these two datasets differ in their acquisition parameters as well as the preprocessing and denoising strategies applied to the data. In the age of multiple large public datasets, the choice of which particular dataset to use in a given analysis is often based largely on convenience and accessibility. Even when researchers make these decisions based on the particular features of each dataset, reasonable arguments can be made about which features should be prioritized, and thus which dataset should be chosen. Ultimately, however, we desire to identify robust effects which replicate across a range of possible datasets and analytic workflows. By attempting to replicate all effects across datasets which differ in these ways, we can provide at least initial evidence of this robustness.

### Timescale estimation

We explored two different methods for estimating timescales. First, we followed the approach introduced by (Murray et al., 2014), in which timescales are estimated based on an exponential fit to the decay of the autocorrelation function of each time series. Second, following (Huang et al., 2018), we characterized timescales based on lag-1 autocorrelation. We found that timescales estimated based on these two methods were very highly correlated in both datasets (r > 0.9), but that lag-1 autocorrelation had significantly higher test-retest reliability than decay-based estimates (see “Evaluation of timescale estimation methods” in the Supplement). These results are consistent with recent work by (Shinn et al., 2021) who found that—for BOLD fMRI time series—lag-1 autocorrelation is highly correlated with estimates of long-range dependence and has higher test-retest reliability than autocorrelation at subsequent lags. Given these results, and following the principle of parsimony (which dictates that given two models with similar performance, we should prefer the simpler model), we chose to focus our analyses on timescales estimated using lag-1 autocorrelation.

We created group-level timescale maps by taking the median of lag-1 autocorrelation values for each region across individuals in each dataset. Because ROI volume can bias timescale estimates by artificially inflating time series autocorrelation (Afyouni et al., 2019; Fallon et al., 2019; Sethi et al., 2017; Shinn et al., 2021), the number of voxels in each ROI was regressed from group-median timescale estimates prior to visualization and during all further analyses.

### Functional connectivity network construction

For each participant, we estimated functional connectivity as the Pearson correlation between the time series of each pair of 246 ROIs defined in the Brainnetome atlas (Fan et al., 2016). We then applied the Fisher r-to-z transform to the correlations estimated from each individual before averaging edges across participants to create a group-level network. Due to uncertainties regarding the interpretation of negative correlations in studies of functional connectivity (Fox et al., 2009; Murphy et al., 2009; Rubinov & Sporns, 2010), all edges with negative z-scores were set to zero. Functional connectivity networks were otherwise un-thresholded. This process was repeated separately for the discovery and validation datasets. In order to identify common functional network communities across datasets, we combined the average FC networks from the discovery and validation data into a single common network prior to community detection. However, measures of regional network topology (i.e. degree, PC, and WD) were computed separately for each dataset based on its respective average FC network.

### Structural connectivity network construction

We constructed a single structural connectivity network from a publicly available, expert-curated voxelwise “population-average” atlas of the human structural connectome (Yeh et al., 2018). This atlas was derived from high-resolution, multi-shell diffusion MRI images from 842 participants from the Human Connectome Project (Van Essen et al., 2013). Complete details of how the atlas was produced can be found in (Yeh et al., 2018), but we review them briefly here. Images for each participant were transformed to MNI space using q-space diffeomorphic registration (Yeh & Tseng, 2011), which preserves spins and fiber geometry and estimates the spin distribution function (SDF) at each voxel. Voxelwise SDFs were averaged across participants to create a group-average template, after which multiple runs of deterministic tractography (Yeh et al., 2013) were used to estimate 550,000 streamlines describing connectivity across the entire brain. All streamlines were then reviewed and labeled by a team of expert neuroanatomists to identify fiber tracts, remove anatomically implausible (i.e. false positive) connections, and manually seed additional streamlines to identify known fascicles which were missing from the initial set of connections. The final atlas contains ∼145,000 voxelwise streamlines describing the representative white matter connectivity of cortical and subcortical regions in the human brain.

Based on this atlas, we estimated structural connectivity as the number of streamlines connecting each pair of ROIs. The resulting whole-brain connectome had an edge density of 0.103, with all but two regions connected to the rest of the network by at least one streamline. However, because the distribution of streamline counts was highly skewed and spanned several orders of magnitude (0 to 1027), we followed the common practice of applying a log transformation to edge weights prior to subsequent analyses (Fornito et al., 2016). Due to the nature of the log transformation (i.e. log(1) = 0), edges with only a single streamline were set to zero, leading to four additional regions becoming disconnected (Fig. S1).

### Multiscale community detection

Prior to calculating community-aware topological measures such as PC and WD, one must first identify (or impose) a “partition” which splits the network into communities and describes which regions belong to each community. While there are many different data-driven methods for identifying communities in complex networks, by far the most common are those which select an optimal partition based on modularity (Q), a mathematically defined quality function which describes the extent to which connections are concentrated within rather than between communities (Newman, 2006). However, all such “modularity maximization” algorithms suffer from a “resolution limit” such that they are unable to detect communities smaller than a given scale which depends on the size of the network and density of within-community connections (Fortunato & Barthélemy, 2007). To counter this limitation, methods have been developed (e.g. (Blondel et al., 2008)) that incorporate a “resolution parameter” (γ) which allows users to adjust the scale at which communities are identified (i.e. fewer larger communities vs. more smaller communities).

However, as noted above, the modular architecture of neural systems manifests at multiple scales, and this is true even for the macro-scale functional and structural brain networks amenable to investigation by human neuroimaging (Betzel & Bassett, 2017). For example, one might initially consider a coarse 2 community partition for human functional brain networks, composed of “task positive” and “task negative” subsystems (Fox et al., 2005). At finer scales, we might identify the “canonical” macroscale functional brain networks (e.g. default, frontoparietal, visual, somatomotor, etc. (Damoiseaux et al., 2006; Power et al., 2011; Smith et al., 2009; Uddin et al., 2019; Yeo et al., 2011)), each of which can in turn be further decomposed into multiple smaller communities (Akiki & Abdallah, 2019; Ashourvan et al., 2019; Meunier et al., 2009; Smith et al., 2009; Yeo et al., 2011). This sort of hierarchical community structure is consistent with a view of the brain as a complex dynamical system which operates across multiple spatial and temporal scales, but poses a challenge to researchers insofar as there is likely no single “correct” scale at which define network communities. Further, because participation coefficient and within-module degree are defined relative to a particular partition, there is no guarantee that these measures will show a consistent relationship across scales to other features of brain structure and function.

To identify network communities at multiple scales, we applied the Louvain algorithm (Blondel et al., 2008) to the structural and functional connectivity networks while varying the resolution parameter (γ) across a range of values (0.5 to 3.5, in increments of 0.1). As γ increases, the algorithm is biased toward detecting smaller and smaller communities, thus also increasing the number of communities identified. Like all community detection methods which rely on modularity maximization, the Louvain algorithm is stochastic, and the partitions it identifies are degenerate (i.e. many possible partitions will have approximately the same modularity) (Good et al., 2010). As such, we ran the algorithm 1,000 times for each value of γ and used consensus-based clustering (Bassett et al., 2013; Lancichinetti & Fortunato, 2012) to identify a single stable partition at each resolution. This resulted in 31 community partitions for each network, with the number of non-singleton communities (# regions > 1) ranging from 3 to 21 for the functional connectivity network, and 2 to 13 for the structural connectivity network.

### Identification of maximally representative community partitions

To identify a maximally representative partition (MRP) for each network, we first computed the variation of information (VI) (Meilă, 2003)—an information-theoretic measure of distance—between each pair of partitions across all 31 scales. We then identified the MRP for each network as the partition with the lowest median VI (and thus the highest similarity with the most other partitions). We note that our approach is conceptually different from how VI (and other measures of partition similarity, such as the Rand index) have most often previously been used to select brain network partitions (i.e. to select the scale at which stochastic runs of a community detection algorithm show the least variability (e.g. (Betzel et al., 2017; Betzel & Bassett, 2017; Shafiei et al., 2019)), and is more closely analogous to the approach taken by (He et al., 2018) and (Bazinet et al., 2021), who used partition similarity to identify domains of stability across scales. Our method builds on this work and is based on the idea that representative features of community structure should be present across a wide range of scales, particularly in a hierarchically organized system like the brain.

For the functional connectivity network, the community partition at γ = 1.8 (8 communities) exhibited the lowest median VI (Fig. S2A), and visual inspection of the pairwise VI matrix (Fig. S2B) confirmed that this partition had good similarity to other partitions across a wide range of scales, ranging from from γ = 1.3 (5 communities) to γ = 3.5 (21 communities). The selected partition at γ = 1.8 also occurs at the visual asymptote of the modularity maximization function (Fig. S2A), such that past this point, modularity (Q) only shows modest gains with increasing resolution. In agreement with prior studies of functional brain organization, the 8 communities identified at this resolution (Fig. S3) show excellent correspondence with “canonical” macro-scale human functional brain networks (Power et al., 2011; Smith et al., 2009; Uddin et al., 2019; Yeo et al., 2011).

For the structural connectivity network, the partition at γ = 2.5 exhibited the lowest median VI (Fig. S4A). However, visual inspection of the pairwise VI matrix (Fig. S4B) revealed that while the partition at this resolution is highly similar to other partitions within the domain of resolutions γ ≥ 2.0, it has relatively low similarity to partitions at coarser scales. In contrast, the partition at γ = 2.0 has only a marginally higher median VI than the global minima at γ = 2.5, while showing good similarity to partitions across a much wider range of resolutions (γ = 0.9 to γ = 3.0, 4 to 12 communities). Further, the partition at γ = 2.0 has the lowest total VI. For these reasons, we judged the partition at γ = 2.0 to be maximally representative. This partition contained 7 non-singleton (N regions > 1) communities (Fig S5).

### Evaluating the relationship between timescales and topological features

The statistical comparison of spatial maps describing different aspects of brain structure and function is a core practice in neuroimaging. However the spatial autocorrelation inherent in brain maps violates the assumptions of independent samples that are key to valid parametric statistical inference (Alexander-Bloch et al., 2018; Bartlett, 1935). This lack of independence inflates the variance of the sampling distribution of the test statistic (e.g. correlations, t-values, etc) and reduces the effective degrees of freedom, leading to inflated Z-scores and high rates of false-positives when using traditional parametric tests. As such, it is necessary to evaluate whether two brain maps show an association *above-and-beyond* that induced by their autocorrelation. To do this, we calculated p-values for all statistical tests using non-parametric permutation testing and 10,000 surrogate timescale maps generated using an autocorrelation-preserving spatial null model (Burt et al., 2020). These were implemented using a custom-developed Python package (https://github.com/danlurie/PyPALM) which facilitates the use of arbitrarily complex null models when evaluating multivariable linear models.

### Correction for multiple comparisons

Given the large number of tests performed in this study, we take a multi-pronged approach to guarding against false-positive results. First, we use the Benjamini-Hochberg method (Benjamini & Hochberg, 1995) to control the False Discovery Rate (FDR) at q = 0.05 across all statistical tests in each dataset (with the exception of a small number of post-hoc tests conducted to evaluate the independence of predictors). We report corrected p-values as p*_perm_^FDR^*, and uncorrected p-values as p*_perm_*. Second, while we report the results from each dataset separately, we require each effect to be significant in both datasets in order to reject the overall null hypothesis.

### Comparing effects and correcting for selective inference

Inference regarding differences in effects necessitates formally testing these differences: it is not enough to simply observe that one effect is significant and the other isn’t (Gelman & Stern, 2006; Nieuwenhuis et al., 2011). The classic example of such comparisons is checking for a significant interaction when conducting an analysis of variance, though our use of multiple models (one for each measure) rather than a single unified model (containing all measures) precludes a formal interaction test. As such, we rely on the method introduced by (Zou, 2007) for comparing correlations. In this approach, a confidence interval is computed for the difference in effects while accounting for dependence (i.e. measures estimated from the same data) and overlapping variables (e.g. X is overlapping in a comparison of corr_X•Y_ and corr_X•Z_). If the confidence interval includes zero, we are unable to reject the null hypothesis for a difference between correlations (and vice versa). Because we are primarily concerned with establishing whether one feature of network topology is more strongly associated with timescale than another (rather than whether the direction of the association differs between features), we computed confidence intervals using the absolute value of partial correlations. However, because we seek to compute confidence intervals for only a portion of the effects we have estimated (i.e. that subset of paired effects for which one effect is significant and the other is not), we are faced with an issue of selective inference (a.k.a circular inference, double-dipping, or “voodoo” statistics (Kriegeskorte et al., 2009; Vul et al., 2009)). Failing to account for such selection risks biased inference. As such, we control the False Coverage-statement Rate (FCR) using the approach proposed by (Benjamini & Yekutieli, 2005) and adapted for neuroimaging by (Rosenblatt & Benjamini, 2014). This method is a generalization of the Benjamini-Hochberg FDR procedure (Benjamini & Hochberg, 1995) to the case of selected multiple CIs. As with correction for multiple testing, we apply the FCR adjustment across all comparisons within each dataset.

### Evaluation of potential confounding due to regional differences in tSNR

We evaluated the possibility that timescale estimates may be confounded by regional differences in signal to noise ratio. For each participant, we calculated tSNR by dividing the temporal mean by the standard deviation of ROI time series from images which had undergone only basic preprocessing (e.g. volume realignment and global intensity normalization). We then calculated the median of these values across participants in each dataset and computed the correlation between timescale and tSNR across cortical ROIs. In the discovery data, we found a weak positive correlation between timescale and tSNR, but this effect was not significant (*r* = 0.175, p*_perm_* = 0.424). In the validation data, the correlation between timescale and tSNR was notably stronger (*r* = 0.374, p*_perm_* = 0.058). This effect was substantially reduced when excluding the 21 cortical areas with the lowest tSNR (i.e. the bottom decile) (*r* = 0.119, p*_perm_* = 0.541). As such, while the main text presents analyses which include all cortical ROIs, we confirmed that all key findings in the validation data were robust to the exclusion of low signal ROIs (see “Robustness to tSNR” in the Supplement). The only effect which failed to replicate in this control analysis was the relationship between timescale and functional connectivity degree, and we note this when reporting the results of that analysis.

## Results

### Mapping cortical timescales with resting BOLD fMRI

Following preprocessing and denoising, we extracted the average BOLD time series from each of 210 cortical regions of interest defined in the Brainnetome atlas (Fan et al., 2016). Timescales are commonly estimated based on the autocorrelation properties of neurophysiological time series (e.g. (Gao et al., 2020; Murray et al., 2014; Raut et al., 2020; Watanabe et al., 2019)), with greater autocorrelation indicating longer timescales. Here, we use the lag-1 autocorrelation, a straightforward measure which we found to be highly correlated with more traditional methods for estimating timescale (i.e. those based on the decay of the autocorrelation function), while having better test-retest reliability than decay-based estimates (see “Evaluation of timescale estimation methods” in the Supplement).

Group-level timescales (Fig. 1) were highly similar between datasets (*r_s_* = 0.79). Cortical areas with the longest timescales (≥80th percentile of all cortical areas in both datasets; Fig. S6) include the dorsal cuneus and precuneus, bilateral superior parietal lobule, and posterior lateral temporal cortex, as well as left anterior medial prefrontal cortex and retrosplenial cortex. Cortical areas with the shortest timescales (≤20th percentile) include bilateral inferior parietal lobule, medial orbitofrontal cortex, anterior cingulate, and portions of medial and inferior temporal cortex, as well as left temporal operculum and anterior insula, and right medial superior frontal gyrus.

**Figure 1:**
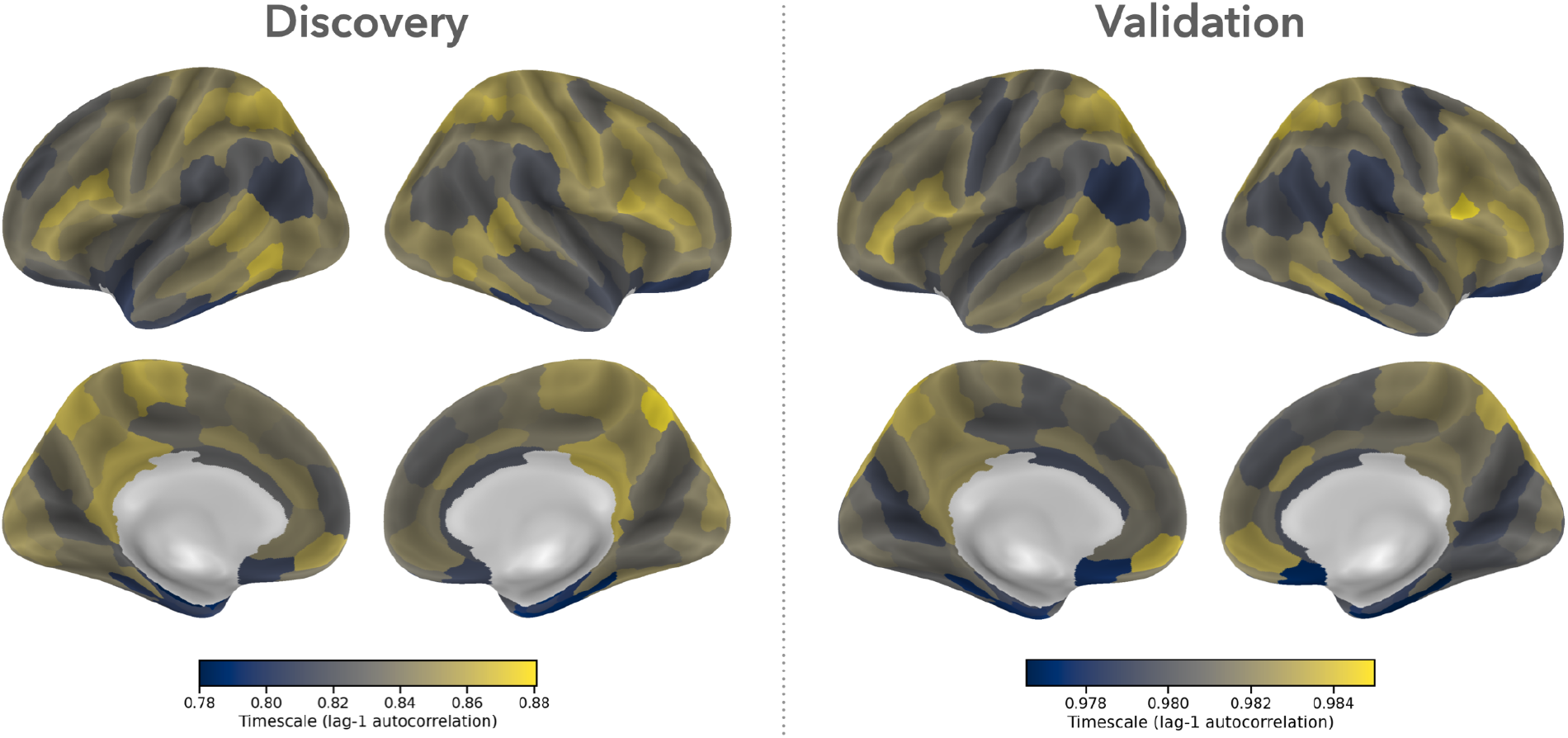
Cortical timescales are consistent across independent datasets. We used lag-1 autocorrelation to estimate the timescale of regional resting BOLD fMRI time series in two datasets of healthy young adults. Our initial (“discovery”) analysis used data from 93 participants from the Nathan Kline Institute-Rockland Sample. We then attempted to replicate our findings in a validation sample of 100 individuals from the Human Connectome Project. The validation data exhibits overall higher values of lag-1 autocorrelation than the discovery data due to the significantly shorter TR (720ms vs. 1400ms).

### Timescales are associated with multiple features of regional brain connectivity

#### Cortical areas with longer timescales tend to have greater structural connectivity degree

The simplest way to characterize and quantify the topology of a brain area is to simply count how many other areas it connects to, and how strong those connections are. In weighted networks (such as those used in this study), this measure is calculated as the sum of the weights of all connections to a region, and is referred to as the “strength” or “weighted degree” of a node. In the interest of brevity—and to avoid confusion when discussing the strength of connectivity between pairs of regions—we refer to this measure simply as “degree”. In mammalian structural brain networks, high degree “hub” regions exhibit preferential connectivity to other highly connected regions, forming a “rich club” of topologically central brain areas whose connections play an important role in integrating information across the network (de Reus & van den Heuvel, 2013; Harriger et al., 2012; van den Heuvel et al., 2012; van den Heuvel & Sporns, 2011, 2013c). High degree hubs are also a key feature of functional brain networks (Achard et al., 2006; Buckner et al., 2009; Cole et al., 2010; Grayson et al., 2014; Tomasi & Volkow, 2011; Zuo et al., 2012). Based on prior investigations into the relationship between network topology and regional dynamics (Baria et al., 2013; Fallon et al., 2020; Gollo et al., 2015; Sethi et al., 2017; Shinn et al., 2021), we predicted that brain areas with higher degree in the functional and structural connectivity networks would exhibit longer timescales.

As predicted, we observed a positive partial correlation (covarying for ROI volume) between timescale and degree in both the functional and structural connectivity networks (Fig. 2). This relationship was significant in both datasets for structural connectivity degree (discovery: *r* = 0.269, p*_perm_^FDR^* = 0.0469; validation: *r* = 0.419, p*_perm_^FDR^* = 0.0006). In contrast, while the correlation between timescale and functional connectivity degree was significant in the validation data when including all cortical ROIs (*r* = 0.385, p*_perm_^FDR^* = 0.0280), this effect disappeared when excluding regions with low tSNR (*r* = 0.124, p*_perm_^FDR^* = 0.544), nor did it reach significance in the discovery data (*r* = 0.300, p*_perm_^FDR^* = 0.1732). These results replicate prior observations of a positive correlation between structural connectivity degree and the timescale of regional BOLD dynamics (Baria et al., 2013; Fallon et al., 2020; Sethi et al., 2017), but fail to replicate reports of an analogous relationship in functional connectivity networks (Baria et al., 2013; Shinn et al., 2021).\

**Figure 2:**
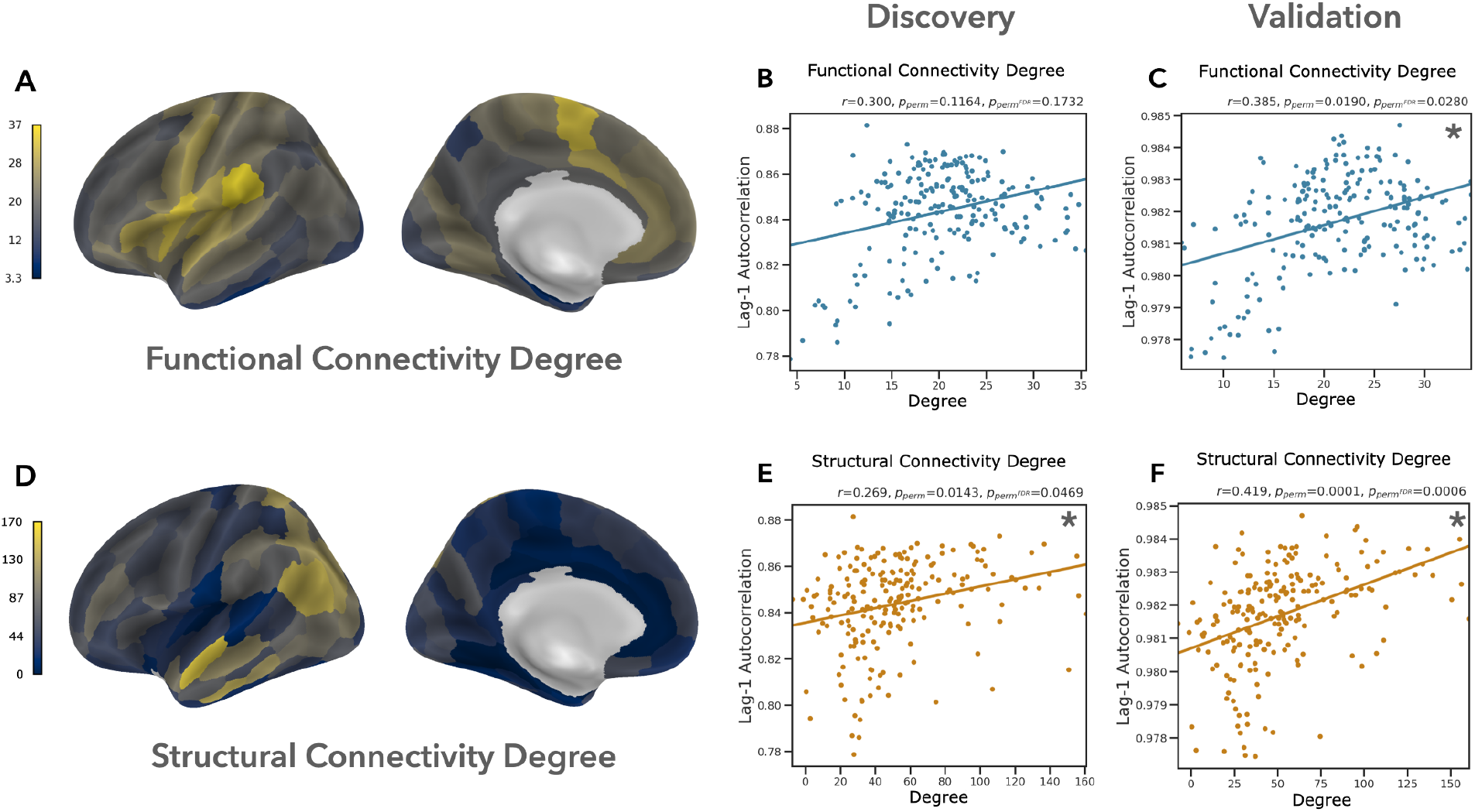
Cortical areas with longer timescales tend to have greater overall structural connectivity. We found that functional connectivity degree (A) was positively correlated with regional timescale, but this effect was not significant in the discovery data (B) or in the validation data when excluding ROIs with low tSNR (Supplement: “Robustness to tSNR”). In contrast, the relationship between timescale and structural connectivity degree (D) was significant in both the discovery (E) and validation datasets (F). Panel (A) visualizes FC degree in the discovery dataset. Asterisks highlight partial correlations which are significant at FDR q ≤ 0.05.

#### The relationship between timescales and the modular structure of human brain networks

In the context of the brain’s modular community structure, a connection between two brain areas can be categorized as being either within-community (if the two brain regions are in the same network community), or between-community (if the two regions are members of different communities). Within-module degree (WD) and participation coefficient (PC) are two complementary measures of regional topology which consider the modular community structure of networks (Guimerà & Nunes Amaral, 2005), and thus provide important additional context about regional connectivity patterns above and beyond that provided by degree alone. In this way, they allow us to take a more nuanced view of the role each region plays in the larger network (Power et al., 2013).

Within-module degree quantifies the extent to which a region has a strong or widespread connectivity to other brain areas within the same network community. However, because communities often differ in size, raw WD values are z-scored across all regions in each community. This is necessary because the preferential within-community connectivity in modular networks will bias regions in larger communities to have a higher degree simply due to community size. Thus the normalized (z-scored) metric provides information about the extent of within-community connectivity of each region relative to other regions in the same community. As such, regions with high WD act as hubs within their respective communities, regardless of the extent to which they exhibit high overall degree. In contrast, participation coefficient provides a measure of the extent to which a region is diversely connected across multiple network communities. A low PC (approaching 0) indicates that a region connects only to other regions within the same network community, while a high value (approaching 1) indicates that connections are distributed evenly across communities. Regions with high PC may act as “connectors” between network communities, and are thought to play an important role in coordinating activity or facilitating the integration or passage of information across communities (Bertolero et al., 2015, 2017, 2018). Given their putative roles in integrating information within (WD) and between (PC) network communities, we predicted that brain areas with high PC and WD would exhibit longer intrinsic timescales. As with degree, we predicted that this relationship would exist in both the structural and functional connectivity networks. These predictions were preregistered (Lurie, 2018).

We took a two-pronged approach to our investigation into the relationship between cortical timescales and PC/WD. First, we identified communities in each network across a wide range of scales and selected a single partition for each network which was most similar to the community structure present across scales. We then computed PC and WD relative to this “maximally representative partition” (MRP) and calculated the correlation between these measures and regional timescales. Second, we evaluated the degree to which our effects replicated when PC and WD were calculated relative to the community structure identified at all other scales. This allowed us to evaluate the robustness of our findings across multiple scales of network organization, as well as the opportunity to identify scale-specific effects (should they exist).

##### Longer timescales are associated with greater within-community functional connectivity across scales, and within-community structural connectivity at coarse scales

As predicted, we found that cortical areas with greater WD in the functional connectivity network tend to have longer timescales (Fig. 3). This relationship was significant in both datasets across the majority of network scales (Fig. 3D), including the scale which we identified as exhibiting the maximally representative community structure (discovery: *r* = 0.368, p*_perm_^FDR^* = 0.0011; validation: *r* = 0.364, p*_perm_^FDR^* = 0.0006; Fig. 3B/C). Regional timescales were also positively correlated with WD in the structural connectivity network. This effect was significant in the validation data across the majority of scales, but only reached significance in the discovery data at the two coarsest scales (Fig. 3H).

**Figure 3:**
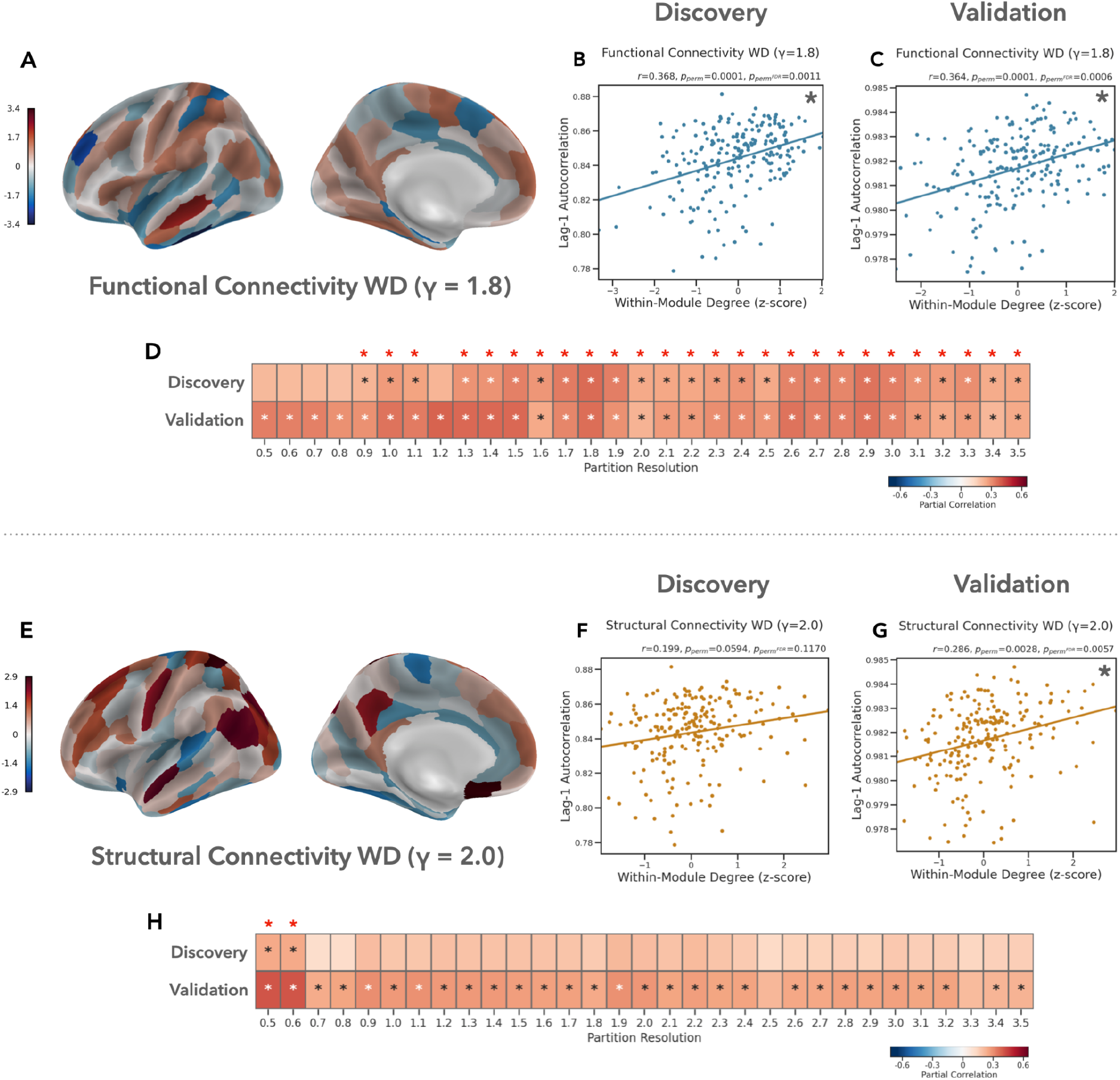
Longer timescales are associated with greater within-community functional connectivity across scales, and greater within-community structural connectivity at coarse scales. (A-D) Brain areas with greater within-community functional connectivity tend to have longer timescales. This relationship was significant in both datasets across the majority of network scales. (E-H) Timescales were also positively correlated with within-community structural connectivity, but this effect only replicated at the two coarsest network scales. Panel (A) visualizes FC WD in the discovery dataset. Asterisks highlight partial correlations that are significant at FDR q ≤ 0.05.

At these coarse scales, cortical areas were split between just two communities (Fig. S7). To investigate the possibility that the relationship between timescale and SC WD was scale-dependent, we compared correlations across scales by computing confidence intervals for the difference in absolute effect size (Zou, 2007). Effects were considered significantly different if the False Coverage-statement Rate (FCR) corrected (Benjamini & Yekutieli, 2005) 95% CI did not include zero. In the validation data, the strength of the relationship between timescale and SC WD was significantly stronger at the two coarsest scales (γ = [0.5, 0.6]) than at finer scales (γ = [1.8, 2.5, 2.6, 2.7, 3.3, 3.4, 3.5]), but there was no significant difference in effect size across scales in the discovery data. These results suggest that hubs of functional network communities tend to have longer timescales across multiple levels of network organization, and that a similar effect exists within structural connectivity networks at very coarse scales.

##### Longer timescales are associated with more diverse structural connectivity

We observed a weak negative correlation between cortical timescales and FC participation coefficient at the majority of network scales. These relationships did not reach significance at any scale in either dataset (Fig. 4A-D). At each network scale, we used confidence interval tests to compare the strength of these correlations to those we observed between timescale and FC WD. In both datasets, we found that the relationship between timescale and FC PC was significantly weaker than the relationship with FC WD at multiple scales (γ = [1.3, 1.4, 1.5, 3.0, 3.1, 3.2, 3.3, 3.4, 3.5]). This suggests that in functional connectivity networks, the extent of within-community connectivity, but not the diversity of between-community connectivity, is associated with longer timescales.

**Figure 4:**
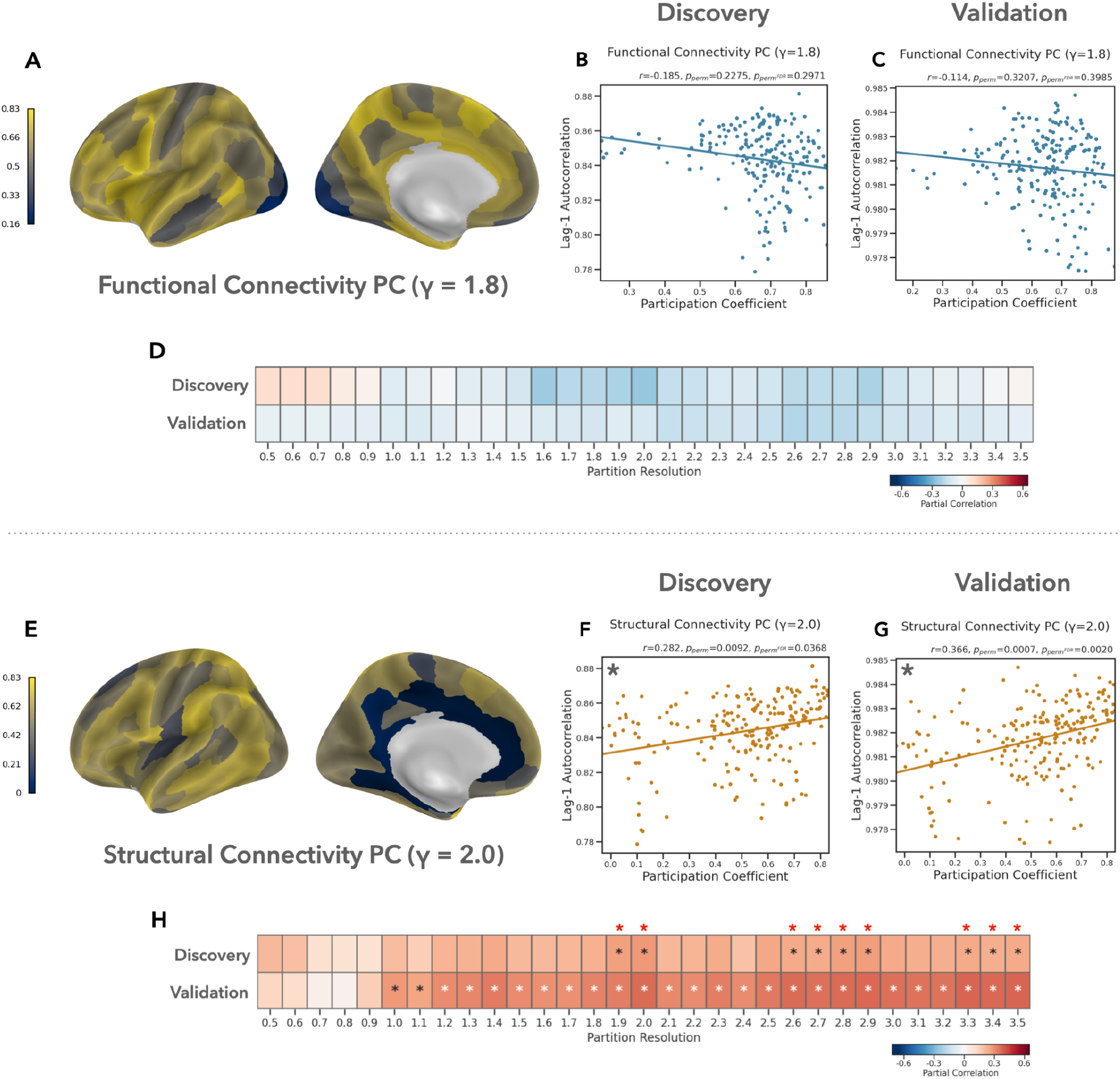
Longer timescales are associated with more diverse structural connectivity. (A-D) In the functional connectivity network, we observed a weak negative correlation between cortical timescales and participation coefficient at most scales, but this effect was not significant at any scale in either dataset. (E-H) In contrast, participation coefficient in the structural connectivity network exhibited a significant positive correlation with cortical timescales in both datasets across multiple scales. Panel (A) visualizes FC PC in the discovery dataset. Asterisks highlight partial correlations that are significant at FDR q ≤ 0.05.

Participation coefficient in the structural connectivity network showed a significant positive correlation with cortical timescales at multiple network scales (Fig. 4E-H), including when PC was computed relative to the maximally representative community structure (discovery: *r* = 0.282, p*_perm_^FDR^* = 0.0368; validation: *r* = 0.366, p*_perm_^FDR^* = 0.0020). In the validation data, this relationship was significantly stronger than the relationship between timescale and FC PC at multiple scales (γ = [1.9, 2.0, 2.9, 3.3, 3.4, 3.5]), but this difference was only significant in the discovery data at the two finest scales (γ = [3.4, 3.5]). Together, these results suggest that longer timescales in the cerebral cortex are associated with structural connectivity which is more diversely distributed across network communities.

##### Functional connectivity WD and structural connectivity PC capture distinct aspects of brain organization

Our observation that cortical timescales exhibit a significant positive correlation with both FC WD and SC PC could in theory result from a correlation between the two measures of regional network topology. To evaluate this possibility, we first calculated the correlation between FC WD and SC PC at each network scale. This revealed that these two measures are only weakly correlated (discovery: mean (SD) = 0.085 (0.048), validation: 0.128 (0.051)). Second, we tested whether the relationship between timescale and SC PC remained significant when controlling for FC WD, and vice versa. For these control analyses, we focused on SC PC and FC WD calculated relative to the maximally representative community partition of each network (γ = 2.0 and γ = 1.8, respectively). The correlation between timescale and SC PC remained significant when controlling for FC WD (discovery: *r* = 0.268, p*_perm_* = 0.0119; validation: *r* = 0.363, p*_perm_* = 0.0019) as did the correlation with FC WD when controlling for SC PC (discovery: *r* = 0.358, p*_perm_* = 0.0009; validation: *r* = 0.361, p*_perm_* = 0.0009). Similarly, we evaluated the possibility that the relationship between timescale and FC WD could be explained by overall functional connectivity degree. We found that the relationship between timescale and functional connectivity WD remained significant in both datasets when controlling for FC degree (discovery: *r* = 0.287, p*_perm_* = 0.0009; validation: *r* = 0.255, p*_perm_* = 0.0099).

### Preferential functional connectivity between regions with similar timescales

Brain areas with similar properties (e.g. in terms of their anatomy (Barbas & Rempel-Clower, 1997; Wei et al., 2019), function (Smith et al., 2009; Yeo et al., 2016), or topology (Bertolero et al., 2017; van den Heuvel & Sporns, 2011)) preferentially connect to each other. Based on this principle of “similar prefers similar” (Goulas et al., 2016), we tested whether regions with similar timescales are more strongly interconnected than regions with dissimilar timescales, as well as if timescales are more similar between pairs of regions in the same network community compared to pairs belonging to different communities. Because brain areas are more likely to connect to other nearby regions compared to more distant areas (Alexander-Bloch et al., 2013; Bullmore & Sporns, 2012; Ercsey-Ravasz et al., 2013), both of these analyses accounted for the potentially confounding effect of Euclidean distance between regions in addition to the spatial autocorrelation of the timescale map. We also tested whether timescales differed between network communities.

To test for preferential connectivity between regions with similar timescales, we first computed the difference in timescale between each pair of cortical areas. We then calculated the partial correlation (covarying for Euclidean distance) between the absolute value of pairwise timescale differences and the strength of functional and structural connectivity between each pair. We found a small but significant negative correlation between pairwise functional connectivity strength and timescale differences in both datasets (Table 1), suggesting that cortical areas tend to exhibit marginally stronger functional connectivity to other areas with similar timescales. Given the positive correlation we observed between timescale and degree, we conducted a control analysis to ensure that preferential connectivity between regions with similar timescales was not a byproduct of the “rich club” phenomenon (i.e. preferential connectivity between high degree hubs). To do this, we regressed degree from timescale estimates prior to calculating pairwise differences. As before, we observed a significant negative correlation in both datasets.

**Table 1:** Preferential connectivity.

| Model | Discovery | Validation |
| --- | --- | --- |
| FC strength | $r = -0.182$<br>**(0.0005)** | $r = -0.151$<br>**(0.0005)** |
| FC strength<br>(adj. degree) | $r = -0.165$<br>**(0.0005)** | $r = -0.159$<br>**(0.0005)** |
| SC strength | $r = -0.027$<br>(0.6143) | $r = -0.100$<br>**(0.0288)** |
| SC strength<br>(adj. degree) | $r = -0.016$<br>(0.6978) | $r = -0.062$<br>(0.1472) |
| SC conn. vs. not | $t = -9.797$<br>(0.2259) | $t = -9.82$<br>(0.1235) |
| SC conn. vs. not<br>(adj. degree) | $t = -8.618$<br>(0.2935) | $t = -9.054$<br>(0.1236) |

Parentheses contain $p_{\text{perm}}^{\text{FDR}}$ .
Significant effects are highlighted in bold.

Because the population structural connectome we analyzed is sparse (i.e. only ∼10% of possible edges are present), we undertook two complementary analyses of preferential connectivity in the SC network. First, we evaluated the correlation between pairwise structural connectivity strength and timescale differences, but limited this analysis to region pairs connected by at least one streamline. We observed a very weak (but significant) relationship between timescale similarity and structural connectivity strength in the validation dataset, but this effect did not replicate in the discovery data or when accounting for SC degree (Table 1). Second, we binarized the structural connectivity network and tested whether the distribution of timescale similarities was different for connected vs. unconnected pairs of brain areas. As with the correlation analysis, we did not find evidence of preferential structural connectivity between areas with similar timescales. When controlling for regional degree, functional connectivity strength had a significantly stronger correlation with timescale similarity than did structural connectivity strength in both datasets (discovery: Δ*r* = -0.149, FCR-adjusted 95% CI = [-0.206, -0.092]; validation: Δ*r* = -0.097, FCR-adjusted 95% CI = [-0.154, -0.041]). Together, these results suggest that there is preferential functional connectivity (but not preferential structural connectivity) between cortical areas with similar timescales.

To test whether timescales were more similar within vs. between communities, we compared the distribution of timescale differences for pairs of regions in the same community to the distribution of differences for pairs where each region belonged to a different community. We repeated this analysis at every network scale. In the validation data, we found that timescales were significantly more similar within vs. between communities of the functional connectivity network at two intermediate network scales (γ = 2.0 and γ = 1.8, both with 7 communities), but neither of these effects were significant in the discovery data (Fig. S8). In the structural connectivity network, we did not find evidence of greater within-community timescale similarity at any scale in either dataset. Finally, we investigated whether timescales differed significantly between network communities by running an ANOVA at each scale. We repeated this analysis for the structural and functional connectivity networks. The main effect of network community failed to reach significance (relative to the spatial null model) at any scale in either dataset (Fig. S9).

## Discussion

In an analysis of resting BOLD fMRI data, we demonstrate that a fundamental property of regional brain dynamics— their autocorrelation-based timescale—is associated with multiple features of macro-scale brain connectivity. By considering the modular organization of neural systems, our results provide new insights into the relationship between connectivity and dynamics in the cerebral cortex. We find that cortical regions with longer timescales tend to have extensive overall structural connectivity, as well as connections which are more diversely distributed across network communities. Longer timescales are also associated with greater within-community functional connectivity, and we show that functional connectivity between brain areas with similar timescales tends to be stronger than functional connectivity between areas with dissimilar timescales. These effects replicate across two different datasets and a wide range of topological scales.

The topography of timescales we observe in the cerebral cortex is largely consistent with the results of previous studies which have mapped timescales in the primate brain using BOLD fMRI (Baria et al., 2013; Fallon et al., 2020; Ito et al., 2020; Manea et al., 2022; Müller et al., 2020; Raut et al., 2020; Shafiei et al., 2020; Shinn et al., 2021; Watanabe et al., 2019; Wengler et al., 2020). That said, our observation of short rather than long timescales in the inferior parietal lobule departs from the results from the majority of previous studies (though we note that there is substantial heterogeneity in this literature). The cortical areas we found to have the longest timescales are all regions whose activity and functional connectivity have been associated with higher-order cognitive functions rather than low-level sensory and motor processes (Smith et al., 2009; Uddin et al., 2019; Yeo et al., 2016). Conversely, however, we did not find that all higher-order cortical areas have long timescales, and we found that many regions which sit near the top of canonical information processing hierarchies (e.g. limbic areas (Mesulam, 1998)) actually exhibit some of the shortest timescales in the cerebral cortex. Similarly, the majority of sensory and motor regions did not have particularly short timescales. As such, our results (and those from numerous other studies (Baria et al., 2013; Fallon et al., 2019; Manea et al., 2022; Shafiei et al., 2020; Shinn et al., 2021; Wengler et al., 2020)) fail to support the view that BOLD timescales are shortest in unimodal sensory areas and longest in heteromodal association regions (though see (Raut et al., 2020) and (Ito et al., 2020)).

Given the sparse spatial sampling found in the majority of studies which used invasive electrophysiology to study neural timescales (typically only a handful of recording sites at most) (Maisson et al., 2021; Murray et al., 2014; Nougaret et al., 2021; Rossi-Pool et al., 2021; Wasmuht et al., 2018), it is difficult to fully determine the extent to which BOLD timescales conform to or depart from much of the broader literature on neural timescales. However, (Watanabe et al., 2019) found very good correspondence between timescales estimated from simultaneously recorded EEG-fMRI, and recent work by (Manea et al., 2022) was able to directly replicate many electrophysiology-based timescale hierarchies using high-resolution fMRI in macaques. However, results from a recent study which mapped timescales across the human cerebral cortex using invasive recordings found both similarities and notable differences compared to maps of BOLD timescales (Gao et al., 2020), suggesting that the relationship between neural timescales and hemodynamic timescales may be spatially heterogeneous. More work is needed to evaluate the correspondence between timescales estimated from these different modalities and to identify factors which may confound studies of hemodynamic timescales (e.g. signal dropout, proximity to draining veins, or regional differences in characteristics of the hemodynamic response function).

Prior work suggests that brain areas with greater inter-regional structural connectivity (i.e. greater degree) tend to exhibit slower BOLD dynamics (Baria et al., 2013; Fallon et al., 2019; Sethi et al., 2017). Our work replicates and extends these findings in important ways. Most significantly, our analyses considered structural connections both within and between cortical hemispheres, as well as between cortical and subcortical regions. In contrast, the studies by (Baria et al., 2013) and (Fallon et al., 2020) both focused exclusively on connections between cortical areas within each hemisphere^1^. In the population-average structural connectome we analyzed here, interhemispheric connections make up almost half (46%) of all edges present in the network, while connections between cortical areas and subcortical regions make up almost a third (29%). As such, failing to consider these connections when analyzing the topology of structural connectivity networks will inherently provide an incomplete picture of brain network organization and the extent to which each region exhibits widespread or diverse connectivity. Further, we believe ours is the first study on the relationship between regional dynamics and structural connectivity degree to consider and correct for the potentially confounding effects of both ROI volume and spatial autocorrelation. These corrections are far more than a formality; the majority of the null effects we report are significant when failing to correct for one or the other of these confounds. Overall, our results demonstrate that this relationship is replicable, statistically robust, and not unique to the special case of intra-hemispheric connectivity between cortical areas. Possible mechanistic explanations for this relationship come from computational models, which suggest that widespread inputs may stabilize the dynamics of high degree regions (Gollo et al., 2015), and that local synaptic integration may act as a low pass filter on these inputs (Baria et al., 2013).

In contrast to our findings for structural connectivity, we did not observe a significant correlation between timescale and functional connectivity degree in either dataset^2^. That said, we caution readers against drawing any strong conclusions based on these results, as the magnitude of the effects we *did* observe are comparable to those for structural connectivity degree. As such, our findings on the relationship between timescale and functional connectivity degree sit in an ambiguous statistical “limbo” (de Hollander et al., 2014); not themselves significant, but not significantly weaker than the significant effects we did observe. We are aware of two previous human resting fMRI studies which found a tendency for regions with slower dynamics to exhibit greater functional connectivity degree (Baria et al., 2013; Shinn et al., 2021). While additional work is needed to determine exactly which factors are driving the discrepancy between our current findings and those in prior reports, we note that our use of weighted degree and consideration of all positive connections differs from the methods used by (Baria et al., 2013) and (Shinn et al., 2021), both of whom estimated degree as the number of connections exceeding a given threshold.

A major contribution of this work is our demonstration that cortical timescales are related to features of network topology which consider the modular organization of structural and functional brain networks. We show that brain areas with long timescales also tend to have higher within-community functional connectivity, and that this effect is at least partially independent of the extent to which these regions also exhibit high overall functional connectivity degree. Within the modular community structure of functional brain networks, these hubs are topologically positioned to facilitate segregated processing by integrating information within functionally specialized subsystems (Bertolero et al., 2018; Sporns, 2013; van den Heuvel & Sporns, 2013b), and our results suggest that this role is reflected in their dynamics. In contrast, we did not find evidence for a relationship between timescales and the extent to which each cortical area exhibits diverse functional connectivity across communities. This finding runs counter to our initial predictions, but is consistent with the fact that regions with diverse connectivity do not necessarily also exhibit widespread connectivity (i.e. regions with high PC do not necessarily also have high within-community or overall degree; (Oldham et al., 2019)). Indeed, we found that FC PC was negatively correlated with FC WD across scales in both datasets (Fig. S10). Together, these results suggest that long timescales may reflect topologically local rather than global integration within functional connectivity networks. Further, they are consistent with the possibility that timescale hierarchies may exist *within* each functionally specialized network module, though further work is needed to directly test this hypothesis.

We also found that regions with longer timescales tend to have more diverse structural connectivity. This finding stands in contrast to our failure to observe an analogous association between timescale and participation coefficient in the functional connectivity network^3^, and underscores the fact that functional connectivity patterns do not necessarily mirror the organization of the underlying structural connectome. The relationship between structural connectivity and functional connectivity is complex and spatially heterogeneous, and many areas with highly correlated brain activity do not have direct anatomical connections (Suárez et al., 2020; Vázquez-Rodríguez et al., 2019; Zamani Esfahlani et al., 2022). Similarly, as evidenced in our results, network communities in the structural connectome do not necessarily correspond to functionally specialized information processing modules. As such, the functional significance of structural connectome topology is not always readily apparent, and must often be discovered iteratively through experimentation rather than intuition. Our results add to this literature. Recent work suggests that regions with diverse structural connectivity have a topology which favors global multi-synaptic interactions (Bazinet et al., 2021). Further, many the regions identified by Bazinet and colleagues as having the strongest preference for topologically global multi-synaptic interactions overlap with areas we found to have the longest timescales, while many of the areas with the strongest preference for topologically local mono-synaptic interactions overlap with regions we identified as having the shortest timescales. This suggests that long timescales in regions with diverse structural connectivity may reflect the integration of signals across a wide range of remote brain areas, including regions to which they exhibit no direct anatomical connections. Our results may also provide insight into recent reports of timescale disruption in psychiatric disorders (Watanabe et al., 2019; Wengler et al., 2020), as regions with diverse structural connectivity are also more likely to exhibit disrupted activity in these conditions (Crossley et al., 2014).

Finally, we found a weak but significant correlation between the strength of functional connectivity between brain areas and the extent to which they exhibit similar timescales. The magnitude of this effect is comparable to that found in a recent report by (Shafiei et al., 2020), who found a positive correlation between functional connectivity strength and the extent to which two areas exhibit similar time series properties. Shafiei and colleagues also found evidence that the properties of resting BOLD time series are more similar between structurally connected regions than unconnected regions, and that time series properties are more similar between regions in the same network community than between regions in different communities. We did not find evidence for either of these effects, nor did we find evidence that timescales differ significantly between network communities. These results suggest that network communities may contain regions with a diversity of timescales, rather than timescales being similar across regions in each community. Additional work is needed to formally evaluate this hypothesis.

This study has at least seven limitations. First, while we analyzed fMRI data from multiple datasets, we compared timescales estimated from those data to a single common structural connectome. As such, our analyses of each dataset are not fully independent. Second, our analysis focused on group-level estimates of timescale and connectivity, and the effects we observe may not fully generalize in individual participants (Fisher et al., 2018). Third, we used only a single parcellation of cortical and subcortical areas, and it is possible that our results might differ when using alternative region definitions or when investigating timescales and connectivity at the level of individual voxels. Fourth, the parcellation we chose did not include any cerebellar regions. Fifth, recent work suggests that the generative null model we used to account for spatial autocorrelation may not exactly control the false positive rate (Markello & Misic, 2021). Sixth, we focused exclusively on communities identified through modularity maximization, and thus did not consider other kinds of network organization such as core-periphery or disassortative community structure (Betzel et al., 2018; Faskowitz et al., 2018). Seventh, we did not account for the potential confounding effect of time series autocorrelation on functional connectivity estimates (Afyouni et al., 2019; Bartlett, 1935). Our findings should be considered in light of these limitations.

## Conclusion

We found that cortical timescales estimated from resting BOLD fMRI are associated with the modular organization of structural and functional brain networks. In particular, we show that longer timescales are associated with diverse structural connectivity and widespread within-community functional connectivity. In addition, we replicate prior observations that regions with long timescales tend to have higher overall structural connectivity, and find evidence for preferential functional connectivity between cortical areas with similar timescales. Additional work is needed to evaluate whether the observed associations are the result of causal influences of network topology on regional dynamics, or if they reflect common principles of cortical organization which shape both local circuit features and patterns of inter-regional connectivity. Our results should also be considered in the context of uncertainties which remain regarding the correspondence of BOLD timescales to underlying neural dynamics. Future studies may benefit from moving beyond the special case of timescales to explore the diversity of ways in which neural time series vary across the brain, and by investigating connectivity-dynamics associations within individual participants. Overall, our results provide new insights into a key feature of functional brain organization—intrinsic regional dynamics—and provide an important empirical link between the literature on cortical timescales and network-centric perspectives which seek to understand the brain as a complex dynamical system.

## Data Availability

All data analyzed in this study are publicly available as part of the Human Connectome Project and Nathan Kline Institute Rockland Sample.

## Ethics approval

This research was conducted under a protocol approved by the Committee for the Protection of Human Subjects at the University of California, Berkeley.

## Footnotes

1 The work by (Sethi et al., 2017) similarly focused on a single hemisphere, but included connections to and from subcortical areas. That said, it should be noted that this study differs from our work and the studies by Baria and Fallon insofar as it analyzed a directed network derived from tract-tracing in mice rather than undirected connectivity estimated from diffusion tractography in humans.

2 This effect was significant in the validation data in our initial analysis, but became substantially weaker and failed to reach significance when excluding cortical ROIs with low tSNR.

3 Though we note that the difference in effects between FC PC and SC PC was only significant in both datasets at the two finest scales.

