## Supplemental information for "Cortical timescales and the modular organization of structural and functional brain networks"

### Supplemental information for "Cortical timescales and the modular organization of structural and functional brain networks", Lurie et al., 2023 (bioRxiv)

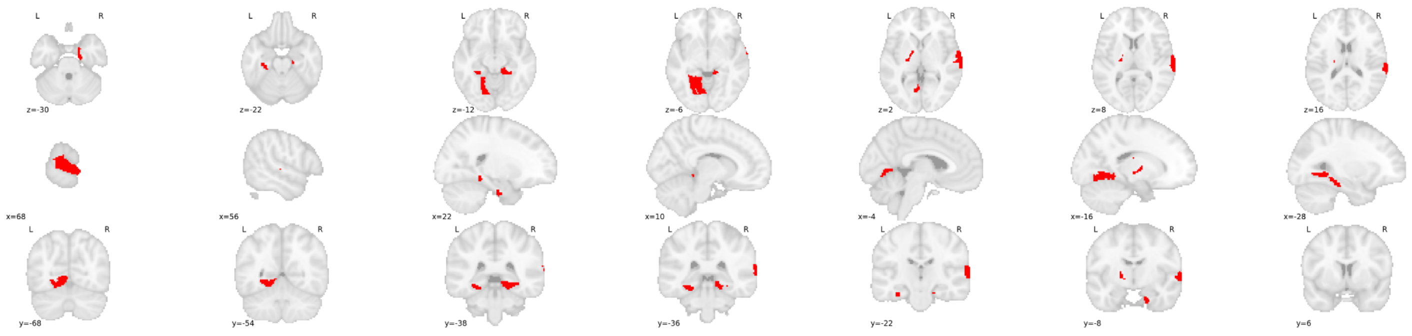


**Figure S1: Regions of interest with no structural connections after log-transforming streamline counts.**

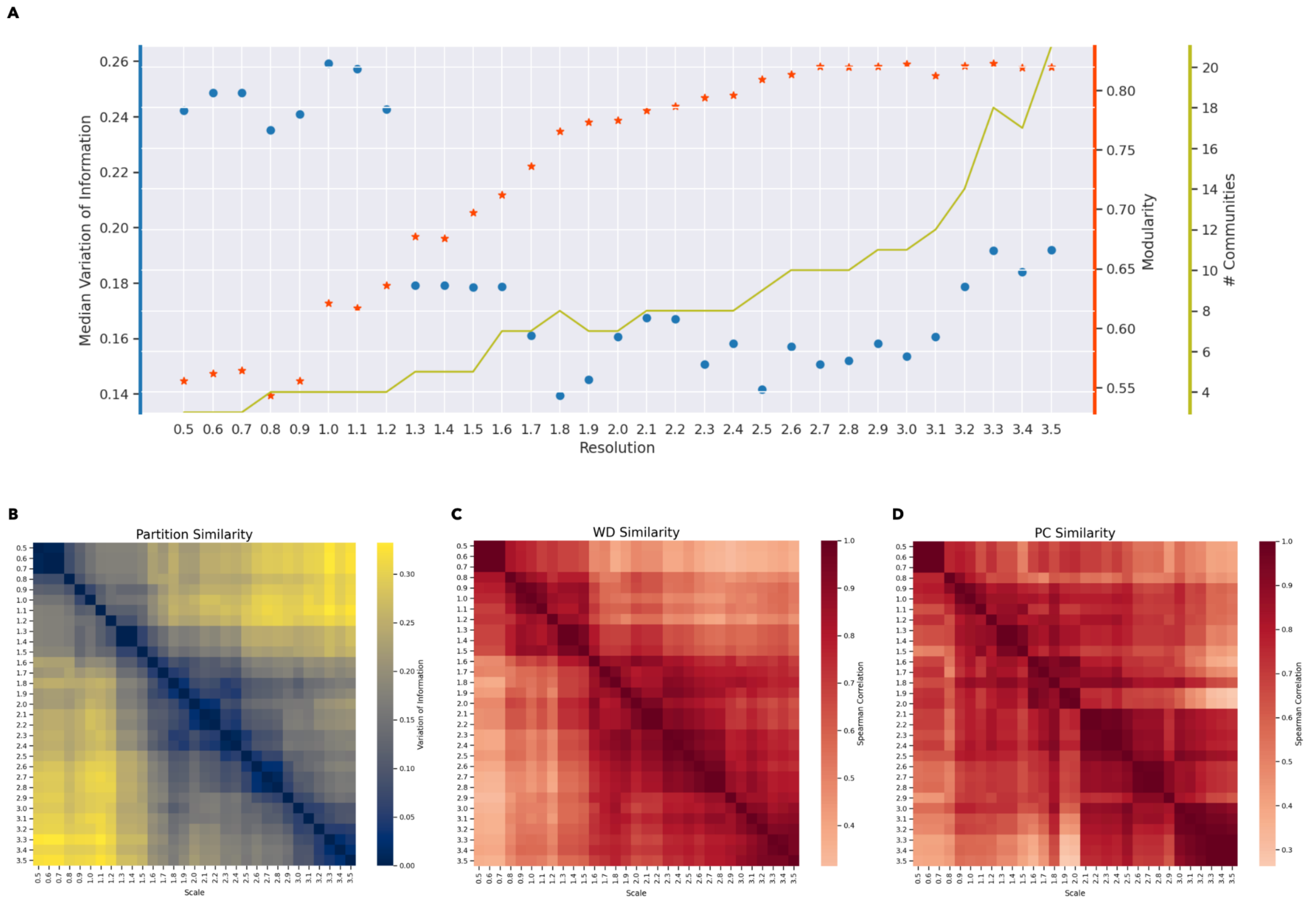


**Figure S2: Multiscale community detection for the functional connectivity network.**

The global minima for median VI was observed at γ = 1.8 (A), leading us to select that partition as being maximally representative of FC community structures across scales. Further support for this selection comes from the fact that this scale occurs at the asymptote of the modularity maximization function (A), and that PC values at this scale are highly similar to those across a wide range of other scales (D).

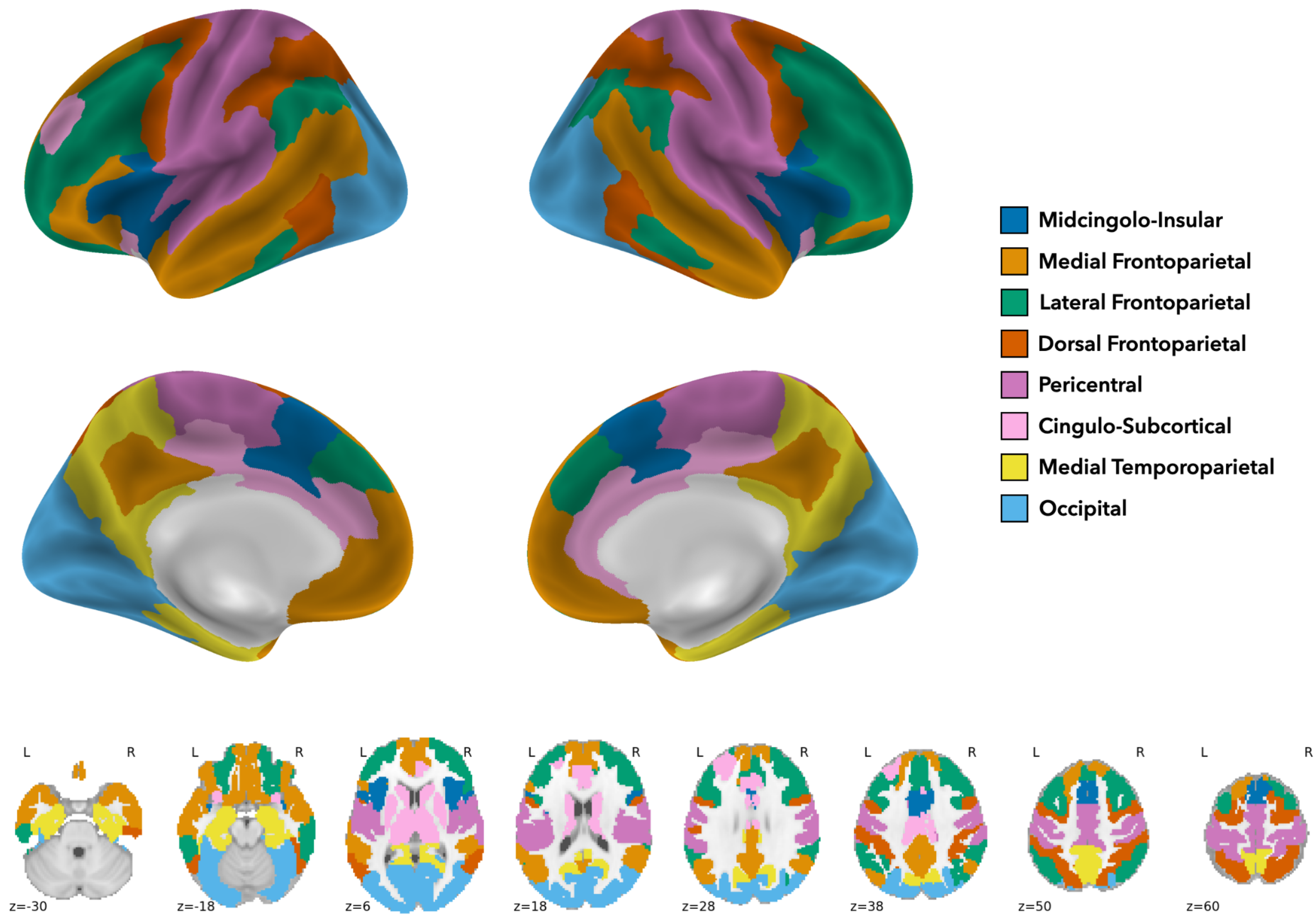


**Figure S3: Maximally representative partition of the functional connectivity network (γ = 1.8, 8 communities).** Community labels are based on the nomenclature proposed in (Uddin et al., 2019).

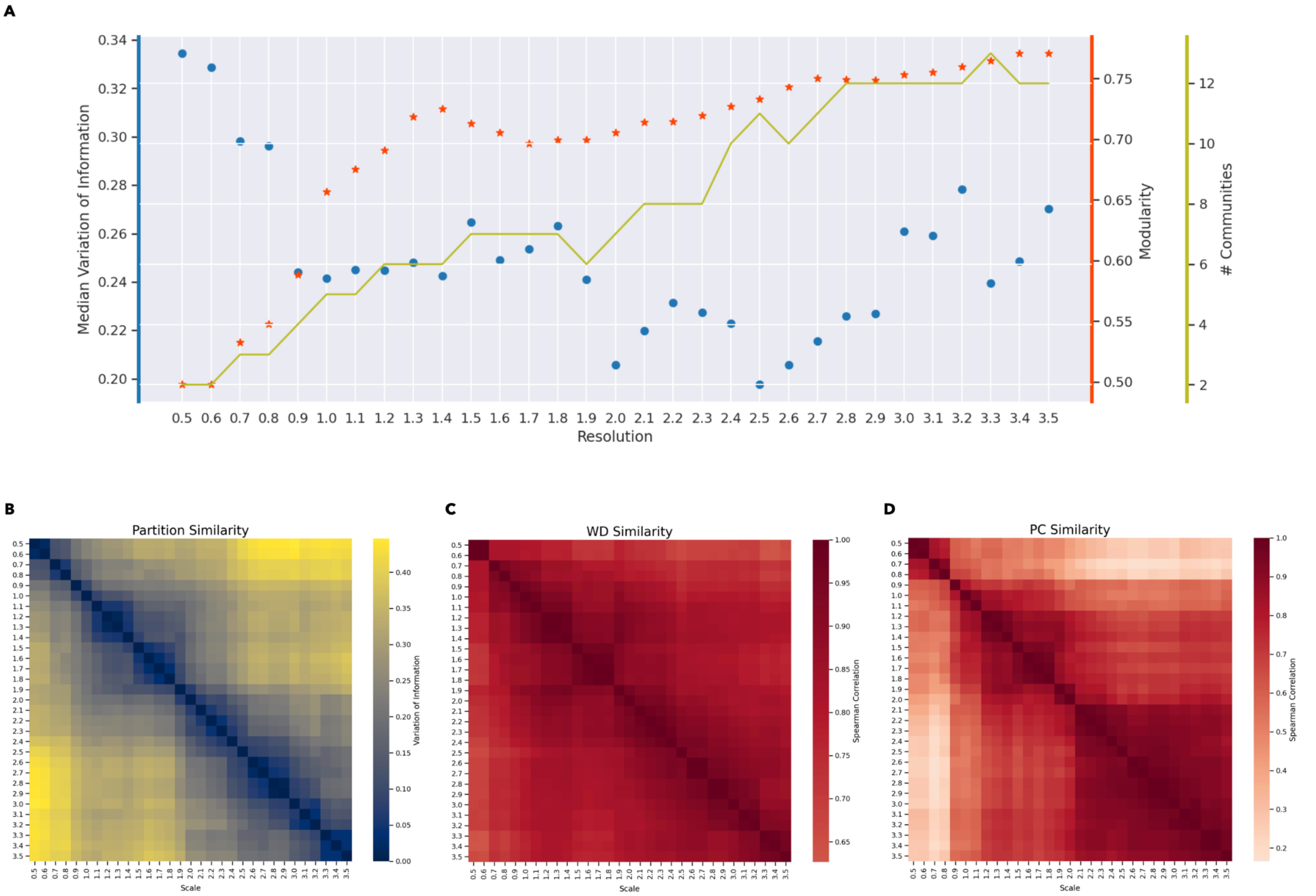


**Figure S4: Multi-scale community detection for the structural connectivity network.**

While the global minima for median VI was observed at γ = 2.5 (A), visual inspection of the partition similarity matrix (B) suggests that the partition at γ = 2.0 showed high similarity with a wider range of other scales while having only a slightly higher median VI. Thus, we selected γ = 2.0 as the scale with the Maximally Representative Partition (MRP).

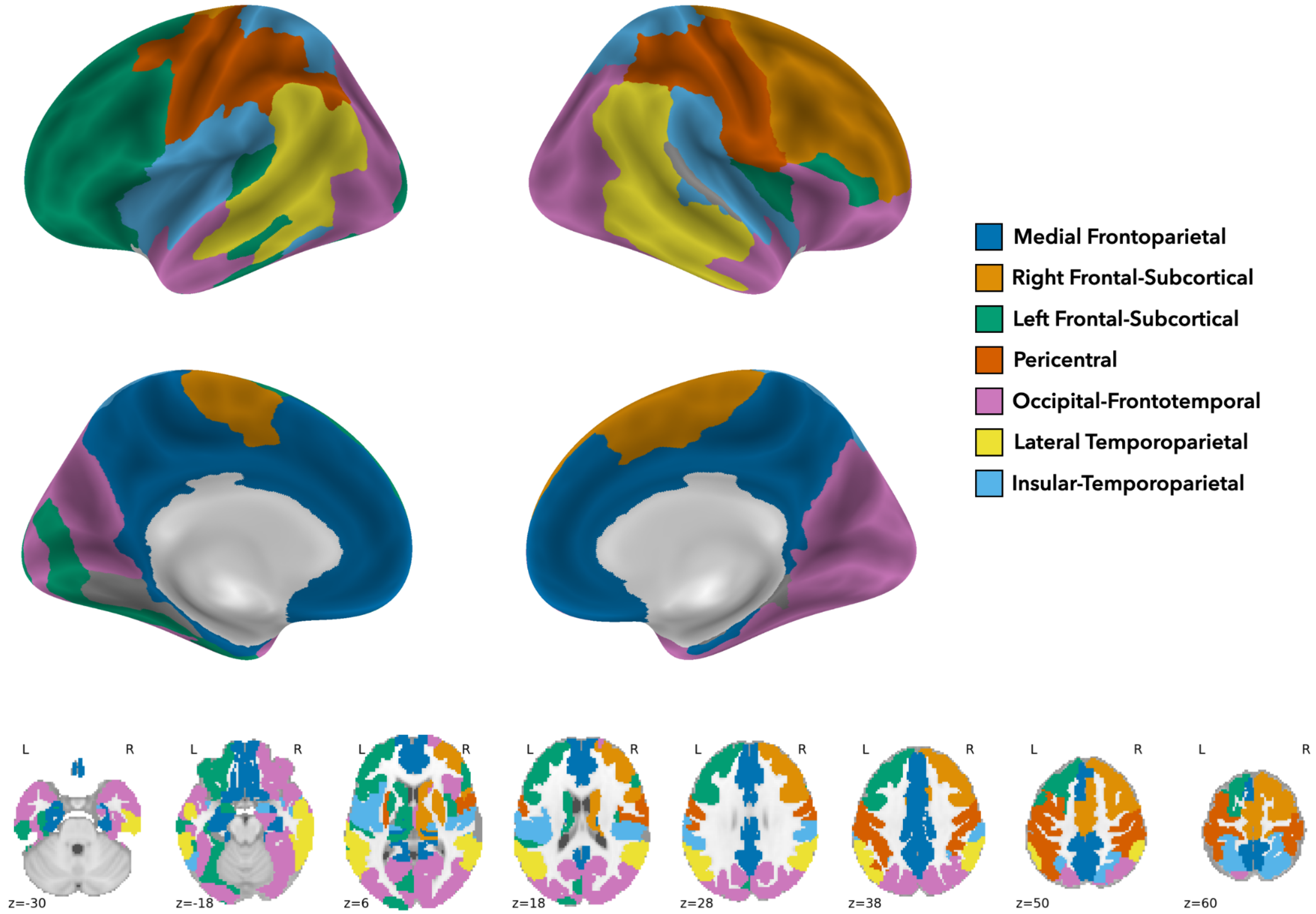


**Figure S5: Maximally representative partition of the structural connectivity network (γ = 2.0; 7 communities).**
Regions assigned to singleton communities (due to disconnection from the larger network) are colored in gray.

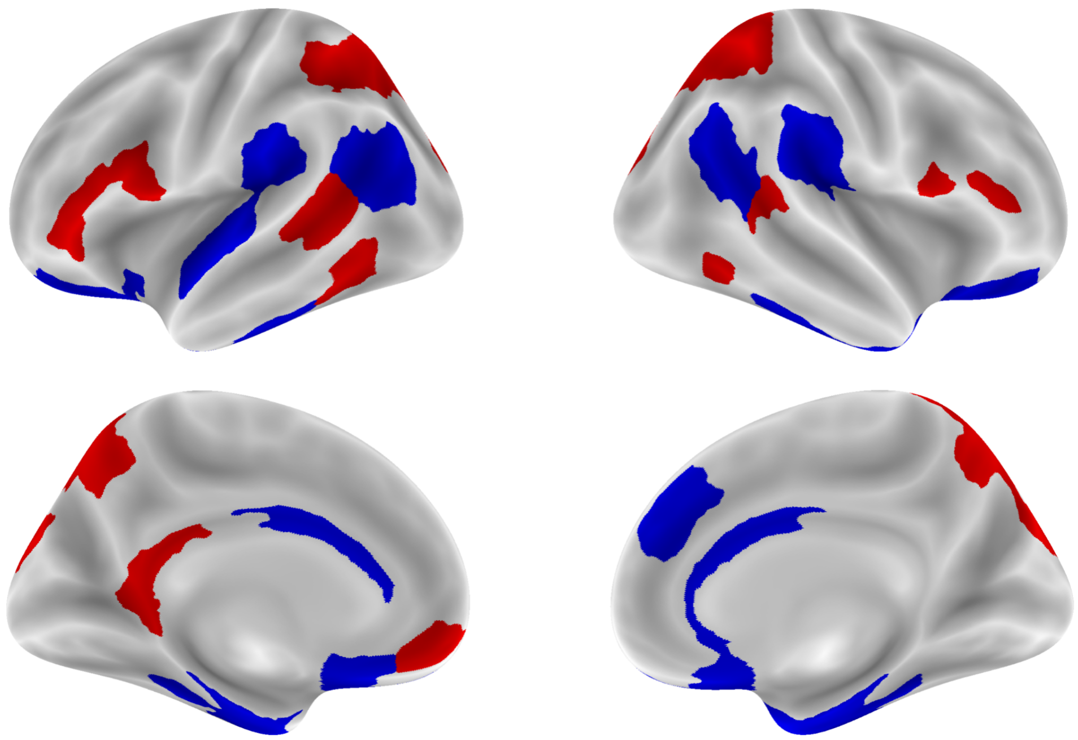

**Figure S6: ROIs with the longest (red) and shortest (blue) timescales.**

ROIs whose lag-1 values fall in the top (red) or bottom (blue) 20th percentile of all cortical areas in both datasets.

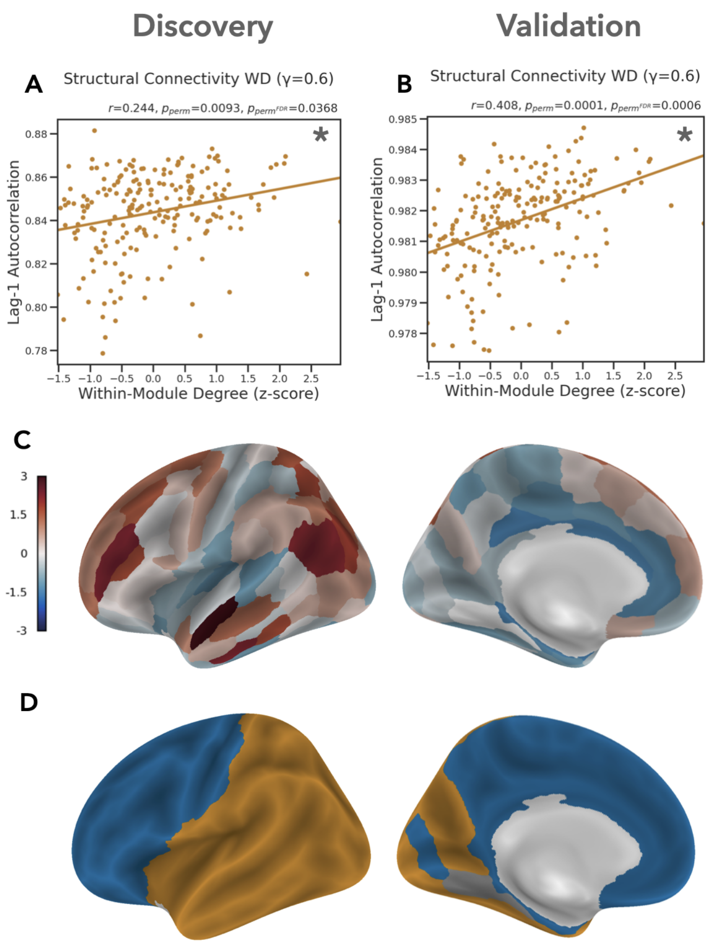


**Figure S7: Correlation between cortical timescales and within-community structural connectivity at coarse scales.**
(A-B) Partial correlation between timescale and SC WD at γ = 0.6. (C-D) SC WD and community structure at γ = 0.6.

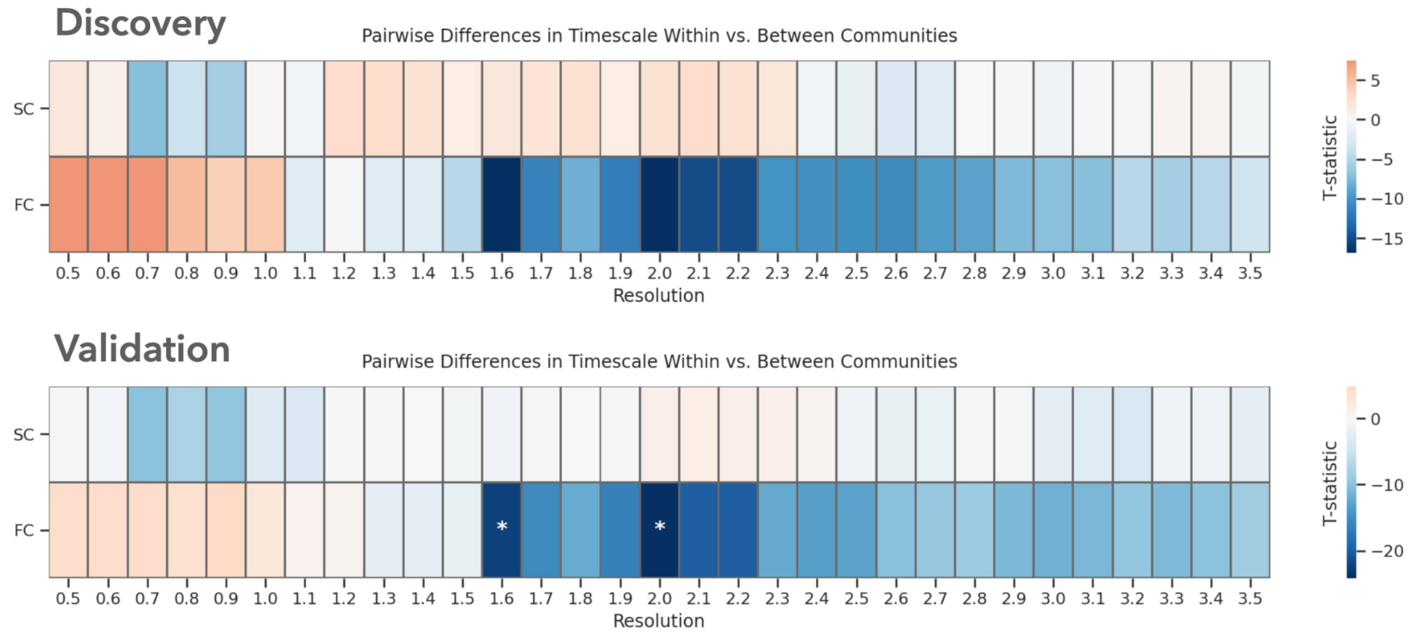


**Figure S8: Similarity of timescales within versus between communities.**

We tested whether timescales were more similar between pairs of regions which belong to the same network community (within) versus pairs which belong to different communities (between). We found significantly greater timescale similarity within versus between functional network communities at two scales in the validation data, but these effects did not replicate in the discovery data. We did not observe any significant effects in the structural connectivity network. Asterisks highlight partial correlations that are significant at FDR $q \leq 0.05$.

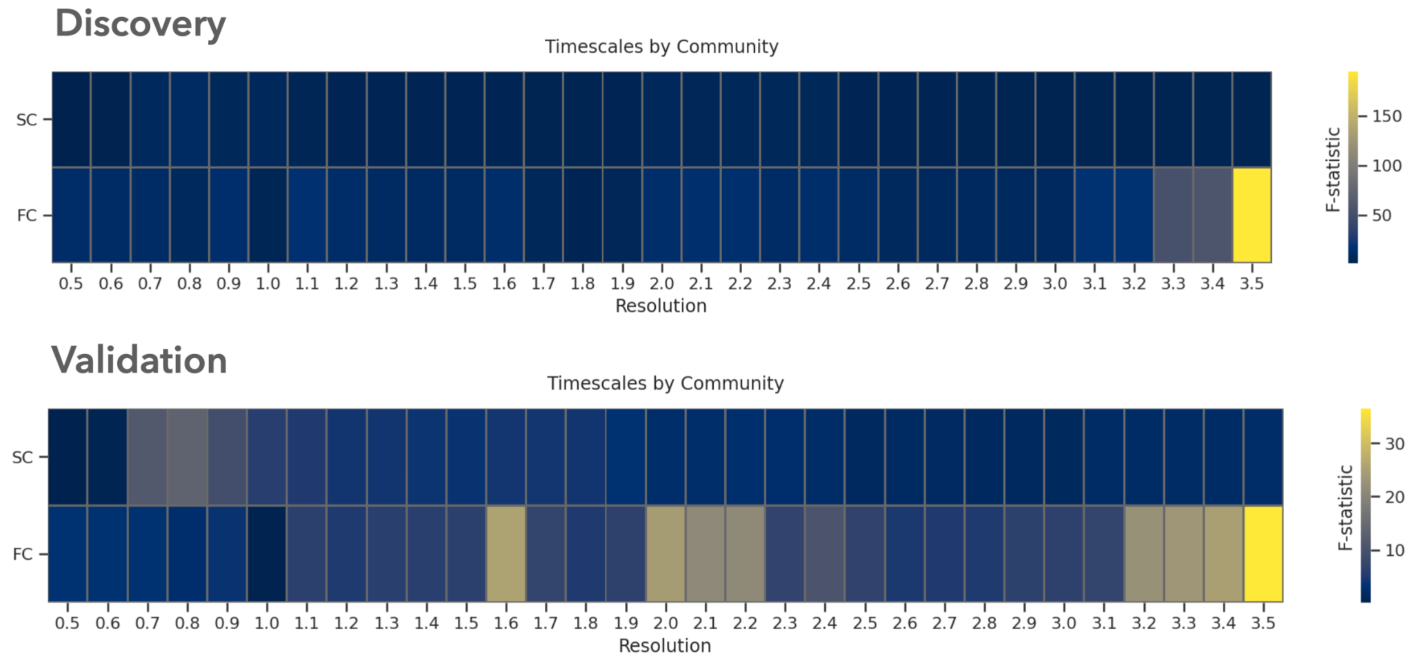


**Figure S9: Differences in timescales between communities.**

We tested whether timescales differed significantly between network communities using an ANOVA. The main effect of community was not significant (relative to the spatial null model) at any scale in either dataset. Asterisks highlight partial correlations that are significant at FDR $q \leq 0.05$.

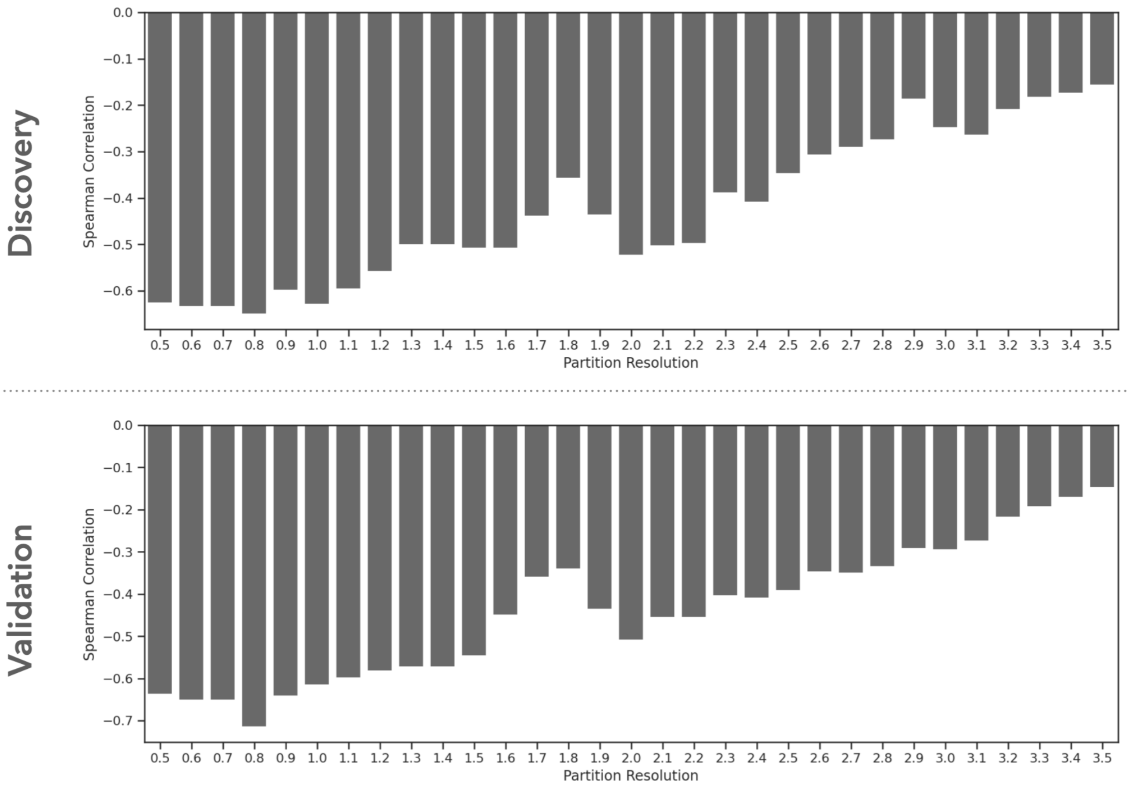


**Figure S10: Correlation between WD and PC in the functional connectivity network.**

#### Evaluation of timescale estimation methods

Following (Murray et al., 2014), we first characterized regional timescales by estimating the exponential decay of the autocorrelation function of each time series:

$$R(k\Delta) \;=\; A\;\left[exp(-\frac{k\Delta}{\tau}) + B\right]$$

Where $\boldsymbol{k\Delta}$ is the lag, $\boldsymbol{A}$ is a scaling constant, $\boldsymbol{\tau}$ is the timescale, and $\boldsymbol{B}$ is an offset constant. We fit this curve at two different maximum lags, the first ("short") selected based on the lag at which autocorrelation had stabilized at or below zero for the majority of regional time series, and the second ("long") as twice the number of lags as the shorter window (Fig. S11A/B). Because the NKI and HCP data have significantly different TRs, the number of lags included for each window differed between datasets. For the NKI data (Discovery), the short window included 10 lags (14 seconds), and the long window included 20 lags (28 seconds). For the HCP data (Validation), the short window included 20 lags (14.4 seconds), and the long window included 40 lags (28.8 seconds). Second, following (Huang et al., 2018), we characterized timescales based on autocorrelation at only the first lag (i.e. the lag-1 autocorrelation). We found that timescale estimates derived from these two methods were highly similar in both datasets (r > 0.9; Fig. S11C/D).

To characterize the test-retest reliability of each method, we additionally analyzed data collected during the second study visit for each participant in the HCP sample, and computed the intraclass correlation of timescale estimates between sessions. We found that timescale estimates based on lag-1 autocorrelation had significantly higher test-retest reliability than those based on the decay of the autocorrelation function (Fig. S11E).

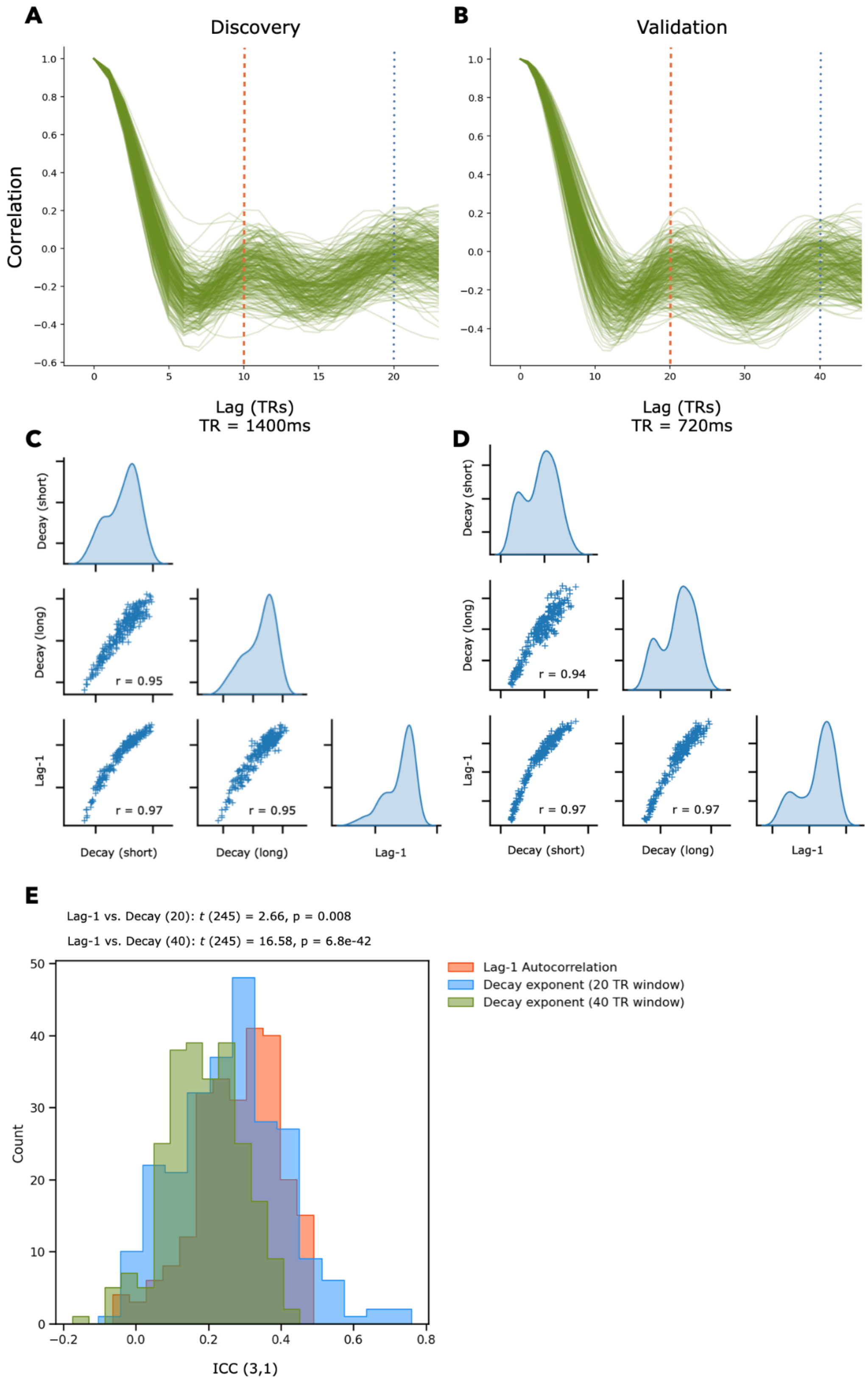


**Figure S11: Comparison of timescale estimation methods.**

(A/B) Autocorrelation functions of all ROIs from two randomly selected participants in each dataset. The orange dashed line and blue dotted line indicate the maximum lag for the short and long decay windows, respectively. (C/D) Distribution of group-median timescales and correlation between different estimates. (E) Test-retest reliability (intraclass correlation) of regional timescale estimates in the validation sample.

#### Robustness to tSNR

To ensure that the observed associations between cortical timescales and network topology were not confounded by tSNR, we repeated our analyses of the validation data after excluding the 21 cortical ROIs (i.e. the bottom decile) with the lowest tSNR. The only effect which was not robust to these exclusions was the correlation between timescale and FC degree, which became substantially weaker ($r$ = 0.124, $p_{perm}^{FDR}$ = 0.544). The relationship between timescale and SC degree remained highly significant ($r$ = 0.433, $p_{perm}^{FDR}$ = 0.003). The relationships between timescale and FC WD, SC WD, and SC PC also remained significant at multiple scales (Fig. S12)

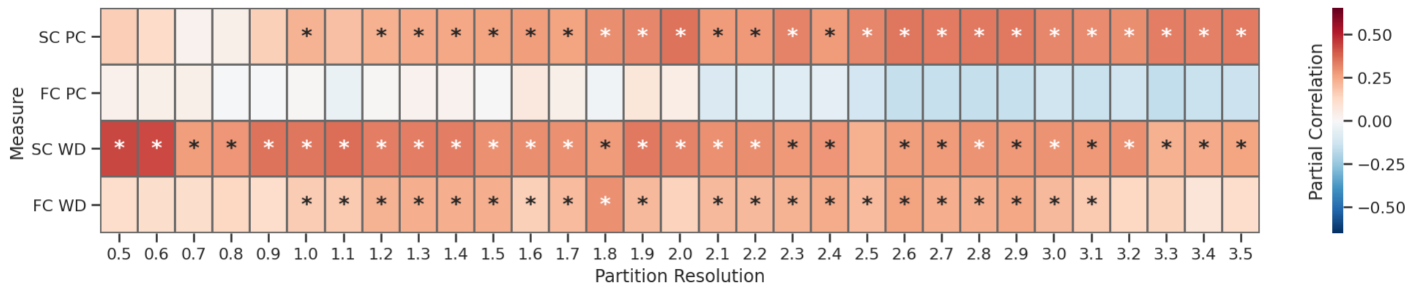


**Figure S12: Replication of topology associations in the validation data after excluding regions with low tSNR .**
Asterisks highlight correlations that are significant at FDR q ≤ 0.05.

#### Deviations from preregistered hypotheses and methods

Here, we briefly describe ways in which the analyses we report here deviate from those specified in our initial preregistration. First, we did not exclude participants in the NKI data based on levels of in-scanner head motion. Second, instead of selecting a single resolution and community partition for each network, we evaluated the association between timescales and topological measures across a wide range of scales. This allowed us to evaluate the robustness of our results. Third, given our finding of significant correlations between regional timescales and topological features (i.e. degree, WD, and PC), we did not undertake the rich/diverse club analyses described in Hypothesis 2 of the preregistration. A modification of the analyses described in Hypothesis 3 of the preregistration are described in a separate forthcoming paper, and as such we do not report them here. Finally, we did not residualize regional timescale estimates relative to age, sex, or head motion prior to combining them across participants.
